# Brief disruption of the microbiome has age-dependent effects on morphine reward and gene expression in the medial prefrontal cortex of adolescent and adult mice

**DOI:** 10.1101/2025.01.02.631117

**Authors:** Rebecca S. Hofford, Jonathon P. Sens, Ava L. Shipman, Violet M. Kimble, Christina Coric, Katherine R. Meckel, Drew D. Kiraly

## Abstract

Adolescence is a critical period for the initiation of problematic drug use, which significantly increases the risk of developing substance use disorders later in life. This heightened vulnerability is partly attributed to the immaturity of the prefrontal cortex, a brain region both essential for decision-making and implicated in drug reward. During adolescence, peripheral systems, such as the gut microbiome, also undergo substantial changes. Emerging evidence suggests that disruptions to the gut microbiome can influence gene expression and drug reward behaviors in rodent models. In this study, we investigated the effects of microbiome disruption on morphine reward and prefrontal cortical gene expression in adolescent and adult mice. Using oral antibiotics to transiently disrupt the microbiome, we found that short-term antibiotic exposure reduced morphine place preference specifically in adolescent mice. In a separate cohort, we observed that antibiotic treatment altered the transcriptomic response to morphine in the medial prefrontal cortex across all age groups. Notably, the transcriptomic changes induced by antibiotics and morphine were age-specific, with distinct gene expression patterns observed in adolescents compared to adults. These findings establish a foundation for future research into the role of the gut microbiome in opioid reward and highlight potential gene pathways underlying age-dependent differences in opioid sensitivity.

## 1. Introduction

Opioid use disorder (OUD) is a major crisis in the United States and around the world. This condition is associated with increased prevalence of chronic health conditions^1,2^ and risk of death from overdose^3,4^. Understanding the biological and environmental factors that contribute to the development of OUD can help identify policies and treatments to help curb opioid use and treat OUD. One predictive factor in diagnosis of a substance use disorder (SUD) is drug use at a young age; age of drug use initiation inversely correlates with risk of developing an SUD later in life^5,6^. For many people, drug use starts during adolescence^7^- a time of life when individuals undergo many physical and social changes. Understanding how adolescents respond to drugs of abuse is necessary to reduce early drug use. Additionally, given the relationship between drug use at young ages and SUD diagnosis later in life, this knowledge could be leveraged to help decrease the total number of SUD diagnoses and overdose deaths.

One contributing factor in the propensity of adolescents to experiment with drugs of abuse is an underdeveloped prefrontal cortex (PFC)^8^. Proper PFC functioning is necessary for healthy working memory and decision-making^9,10^, but this region has also been implicated in drug addiction. Dysfunction in this region has been observed in patients with SUD^11^, and it has been proposed that individuals with pre-existing abnormalities in PFC function are more prone to problematic drug use^11^. While the rodent frontal cortex is not as complex as humans’, the rodent PFC also contributes to decision-making and drug reward^12–14^ and is rapidly developing during adolescence^15–17^.

While the PFC of adolescents is undergoing many changes, the physiological effects of adolescence are not restricted to the brain. Puberty occurs during adolescence, as well as changes in body composition and rapid growth during the teenage years. Partially due to changing energy needs and diets, the adolescent gut microbiome undergoes drastic shifts in humans^18,19^ and is undergoing changes between the juvenile period and adulthood in rodents^20,21^. Increasing evidence suggests that the gut microbiome has far reaching impacts on health and disease, including disorders of the central nervous system. In fact, causal studies in animals have shown that changes to the microbiome have drastic impacts on outcomes in models of neurodegenerative^22^ and psychiatric disorders^23–28^. While most studies are conducted in adulthood, alterations to the microbiome in adolescence also show effects on anxiety-like behaviors^23^ that last into adulthood^29^ and produce alterations to social behavior after traumatic brain injury^30^. Additionally, one week of Abx treatment during early postnatal life, during pre-weaning, or immediately post-weaning produced differing effects on behavior and gene expression in the brain during adulthood^31^, demonstrating that the behavioral and molecular effects of microbiome manipulation differ depending on age of microbiome disruption.

In addition to measures of anxiety and depression, prior work has shown that the microbiome can influence drug reward and reinforcement^32,33^. Our lab and others have found that manipulation of the microbiome alters cellular and behavioral responses to cocaine^34–36^ and opioids^37–40^. We have shown that knockdown of the microbiome with oral antibiotics (Abx) decreases morphine conditioned place preference (CPP)^38^ and increases self-administration of low-dose fentanyl and increases breakpoint during progressive ratio^37^ in adults.

Additionally, there are several studies investigating how Abx alters the molecular and cellular response to drugs of abuse^36–39^. In studies employing sequencing or proteomic analyses, more differentially expressed genes (DEG) or proteins were found in animals with a reduced microbiome and receiving cocaine^36^ or opioids^37,38^. Additionally, the functional role of the DEGs from microbiome-depleted animals is different than those from microbiome-intact animals, with more upregulated genes related to histone and chromatin modification found after microbiome knockdown. These studies have largely focused on the nucleus accumbens (NAc), a brain region strongly implicated in drug reward and reinforcement. However, other studies suggest that the mPFC is also sensitive to changes to the gut microbiome^30,41,42^. Given the importance of the mPFC in drug reward^12–14^, and the fact that the mPFC is changing more dramatically than the NAc during adolescence^15^, the current study measures morphine reward after microbiome manipulation with Abx and explores the molecular consequences of morphine in the mPFC with and without oral Abx. Unlike our prior work that employed 14 days Abx treatment before behavior, a shorter exposure period of Abx is utilized to restrict microbiome disruption and morphine exposure to within the early adolescent period (starts during the juvenile period PND25 and ends PND35) and compares results to those found in young adults (starts PND55 and ends PND65). Data from this study explores how brief Abx exposure alters morphine reward and the mPFC transcriptome in adolescents and adults.

## 2. Materials and Methods

### 2.1. Animals

Male and female adolescent (PND 25 at start of drink treatment) and adult (PND 55 at start of drink treatment) C57BL/6J mice (Jackson Laboratories, Bar Harbor, ME, USA) were used for all experiments. Mice were housed 4 or 5/cage with age- and sex-matched animals in a humidity and temperature-controlled vivarium on a 12h light:dark cycle. Mice were left undisturbed in the colony for four to seven days before the start of drink solutions. At age PND 25 or PND 55, cages were randomly assigned to control H_2_O or antibiotics (Abx) treatment. All animal procedures were approved by the IACUC at Mount Sinai School of Medicine and all animal procedures conformed to ARRIVE guidelines. Separate groups of mice were used for conditioned place preference and sequencing experiments.

### 2.2. Drugs

Morphine sulfate pentahydrate was provided by NIDA drug supply program and was dissolved in isotonic saline. Morphine and saline were injected subcutaneously at a volume of 10ml/kg.

### 2.3. Drink solutions

Mice were placed on control H_2_O or a cocktail of antibiotics on PND 25 or PND 55. Antibiotics consisted of 0.5 mg/ml vancomycin (Chem-Impex International), 2 mg/ml neomycin (Fisher Scientific), 0.5 mg/ml bacitracin (Research Products International), and 1.2 microg/ml pimaricin (Infodine Chemical) dissolved in drinking water^36–38^. Mice remained on drink solutions for five days before the start of behavior or injections and all mice remained on their drink solutions throughout the entire study.

### 2.4. Conditioned place preference

Adolescent and adult male and female mice (*n* = 11-14, 5-8/sex) on H_2_O or Abx underwent conditioned place preference (CPP) for morphine using Med Associates boxes and software as previously described^35,38^. The testing apparatus consisted of three distinct chambers-two larger chambers to either side of a small middle chamber. The end chambers contained unique wall patterns and floor textures to distinguish them from each other (one chamber had gray walls and a small grid floor and the other chamber had black and white striped walls and a larger grid floor). The entire CPP task took place over 5 consecutive days: one day of pre-test, three conditioning days, and one test day. For pre-test day, mice were placed into the small middle chamber and were allowed to explore all three chambers for 30 minutes. Morphine chamber was assigned such that there was no baseline preference within any group. On conditioning days, mice received injections of saline (s.c.) and were confined to one end chamber for 60 mins in the morning (saline chamber). No less than 3 hours later, mice were injected with 5.6 or 10 mg/kg morphine (s.c.) and were confined to the opposite end chamber for 60 mins (morphine-paired chamber). On test day, mice were again allowed to explore all three chambers of the CPP apparatus for 30 mins. Mouse location during pre-test and test sessions were recorded by laser beam breaks. CPP score was calculated as: [(time in morphine-paired chamber on test day) – (time spent in saline chamber on test day)].

### 2.5. 16S sequencing

Separate groups of adolescent and adult male and female mice on H_2_O or Abx were given once daily injections of saline or 15 mg/kg morphine (s.c.) for 5 days (*n* = 5 for all groups). Twenty-four hours after their last injection, mice were rapidly decapitated. Cecal content was collected and flash frozen on dry ice. DNA was isolated from cecal contents using DNeasy PowerSoil kit (Qiagen) via kit instructions with the additional bead beating step. DNA concentration was quantified using a NanoDrop1000. PCR amplification was achieved using primers (341F/805R) targeting the variable V3 and V4 region of the 16S rRNA bacterial genome. DNA fragments were sequenced on an Illumina NovaSeq (2x 250 bp). Using Divisive Amplicon Denoising Algorithm 2^43^, data were filtered, dereplicated, and paired ends were merged. Chao1 and Simpson Indices were used to assess alpha diversity. Principle coordinates analysis plots were generated using the unweighted Unifrac distance as an assessment of beta diversity using QIIME2 software^44^. Observed taxonomic units (OTU) were identified by comparing their genetic sequences to reference bacterial genomes using SILVA (Release 132)^45^ with confidence set to 0.7. Male and female data were sequenced and analyzed separately.

### 2.6. RNA sequencing

#### 2.6.1. Tissue collection and RNA extraction

The same mice used for 16S sequencing were used for RNA sequencing (*n* = 5 for all groups). Twenty-four hours after their last injection, all mice were euthanized as described above. Whole brain was removed and sectioned using a 1mm partitioned brain mold. The dorsal mPFC, containing prelimbic and anterior cingulate cortices, was identified in the rostral most 4mm of brain dorsal and medial to the corpus callosum. A 12g blunt needle was used to remove a 2mm thick section of mPFC. Tissue punches were flash frozen on dry ice and stored at -80° C. RNA was extracted using RNeasy Mini kit (Qiagen) via kit instructions.

#### 2.6.2. Sample preparation and sequencing

Libraries were prepared using the Fast RNA-seq Library Prep Kit V2 (abclonal Technology). Fragments were read to 150 bp (paired-end) on an Illumina HiSeq2500 to an average depth of 25M reads. Reads were removed if they contained adaptor contamination, when nucleotides were uncertain for 10% or more of the read, or the base quality was less than 5 for 50% of the read. Whole samples were excluded from analysis if the difference between their adjusted number of read counts and the average number of read counts were >80% greater or less than the range of the remaining samples’ read counts. Two adult male H_2_O-Sal samples were excluded from further analysis due to low number of reads and two male samples (one adolescent H_2_O-Sal and one adolescent Abx-Mor) and two female samples (two adolescent H_2_O-Mor) were excluded for having too many reads. Male and female samples were sequenced and analyzed separately.

#### 2.6.3. RNA-sequencing bioinformatics

Reads were aligned to the Mus musculus 10 reference genome using the aligner STAR v2.7.10a^46^. Aligned reads were normalized using DESeq2^47^ and differential expression analysis was conducted with ExpressAnalyst^48^ software. Pairwise comparisons of all genes were conducted between adolescent and adult controls (H_2_O-Sal) to determine age-specific gene expression signatures. Additional comparisons were conducted between each treatment group (H_2_O-Mor and Abx-Mor) and their age- and sex-matched control (H_2_O-Sal) to determine the effects of each treatment within each age and sex. Genes were considered differentially expressed if they met a statistical significance of *p* < 0.05 after FDR correction. Significantly up- and downregulated gene lists for each pairwise comparison were separately uploaded to G:profiler^49^ and significant pathways were identified using an FDR corrected *p* < 0.05.

To generate heatmaps of the adolescent signature genes, the log 2 fold change of raw counts from genes significantly differentially regulated between adolescent and adult controls (H_2_O-Sal) were rank ordered from largest downregulated to largest upregulated using the website Morpheus (Broad Institute, https://software.broadinstitute.org/morpheus). Next, the log 2 fold change in expression of the adolescent-specific genes were calculated between the adolescent treatment groups (H_2_O-Mor and Abx-Mor) and adult controls (H_2_O-Sal). Gene expression information from treatment groups were aligned to the adolescent control group to visualize overall alterations in the developmental gene signature across treatments.

For pathway term heatmaps, pathway term titles were searched for select words of interest and grouped by category for each comparison. For the “development” category, pathway terms containing the words “development” or “genesis” were counted. The “vascular” category counted pathways with the prefix “vascul-“, or “angio-“, or the word “vessel”, the “histone modification / transcription” category counted “histone”, “chromatin”, “DNA”, or “transcription”, the “immune” category counted “immune”, “cytokine”, or “microglia”, and the “extracellular matrix” counted “extracellular matrix” or “collagen”. If a pathway term contained more than one keyword, it was only counted once for that category. The number of unique terms in each category were counted and normalized to total term number for each comparison.

Finally, weighted gene co-expression network analysis (WGCNA) was performed as done previously^36,50^ using the WGCNA R package^51^. WGCNA was performed with individual female sample weights first determined with the ‘signed hybrid’ network (i.e. negatively correlated genes are assumed not connected). Signed adjacency matrices were generated using a soft-power parameter via a scale-free topological fit to at least a power of 0.8 whilst maximizing mean connectivity. From the determined scale of independence and mean connectivity calculated, we used a soft thresholding power of 7. Subsequently, a topological overlap matrix (TOM) was generated through the adjacency matrix to perform hierarchical clustering. To identify unique modules, a one-step network was constructed via blockwise modules with unsigned topological overlap matrices. Distinct modules were identified and labeled using the Dynamic Tree Cut method. Notably, only networks with a minimal network size of > 40 genes were used for subsequent analysis.

To determine relationships between modules and treatments, Pearson correlation coefficients between module eigengenes or treatment status were calculated. Importantly, heatmaps and pathway analyses for WGCNA figures included all genes assigned to a specific module (i.e. eigengenes with scores less than 0.99 were included). Modules were chosen for further analysis if they were significant in at least one treatment group, with priority given to modules showing significant positive correlations in Abx-Mor groups. Of the 15 modules meeting this criterion in adolescent Abx-Mor females, we chose one module that also had a significantly negative correlation in adolescent H_2_O-Mor group (black) and another module that had the largest number of associated pathways (blue). Additionally, of the seven modules that had a significant positive correlation in adult Abx-Mor only, we chose to examine the pathway with the largest number of associated pathways (light green). For these three modules, the top 25 driver genes or “nodes” within these modules were identified using cytoHubba’s Maximal Clique Centrality (MCC) feature within Cytoscape^52^, which measures the relatedness or connectedness of all the nodes in the module. However, all nodes from these modules were included in pathway analysis via G:profiler^49^.

### 2.7. Statistics

Conditioned place preference score for adolescents and adults were analyzed separately using two-way ANOVAs with drink and morphine dose as between-subjects factors. *A priori* planned comparisons were conducted between H_2_O and Abx mice at each morphine dose using Sidak’s multiple comparisons. This study was not powered to detect sex differences in behavior, so males and females were analyzed together.

The alpha diversity metrics of Chao1 and Simpson were analyzed using two-way ANOVAs. *A priori* pairwise comparisons were conducted between the control samples (H_2_O-Sal) and each treatment group (H_2_O-Mor and Abx-Mor) within each age using Sidak’s multiple comparisons.

## 3. Results

### 3.1. Brief microbiome disruption reduces morphine place preference in adolescents but not adults

To determine the effects of brief Abx exposure on morphine reward in adolescents and adults, mice underwent morphine CPP after five days of Abx or H_2_O. Pre-adolescent (PND 25) and adult (PND 55) male and female mice were given Abx in their drinking H_2_O or remained on control H_2_O for five days before CPP (**Fig. 1A**). Interestingly, there was a main effect of Abx reducing adolescent (*F*_(1,45)_ = 6.277, *p* = 0.0159), but not adult (*F*_(1,49)_ = 0.3898, *p* = 0.5353) morphine place preference (**Fig. 1B-C**). Planned comparisons indicated a significant difference between adolescent mice treated with H_2_O and Abx at the 10 mg/kg dose (*p* = 0.0247). There were no significant pairwise comparisons in adult mice, only a main effect of morphine dose (*F*_(1,49)_ = 8.898, *p* = 0.0044) that was not present in adolescents (*F*_(1,45)_ = 0.6184, *p* = 0.4358).

**Fig. 1.**
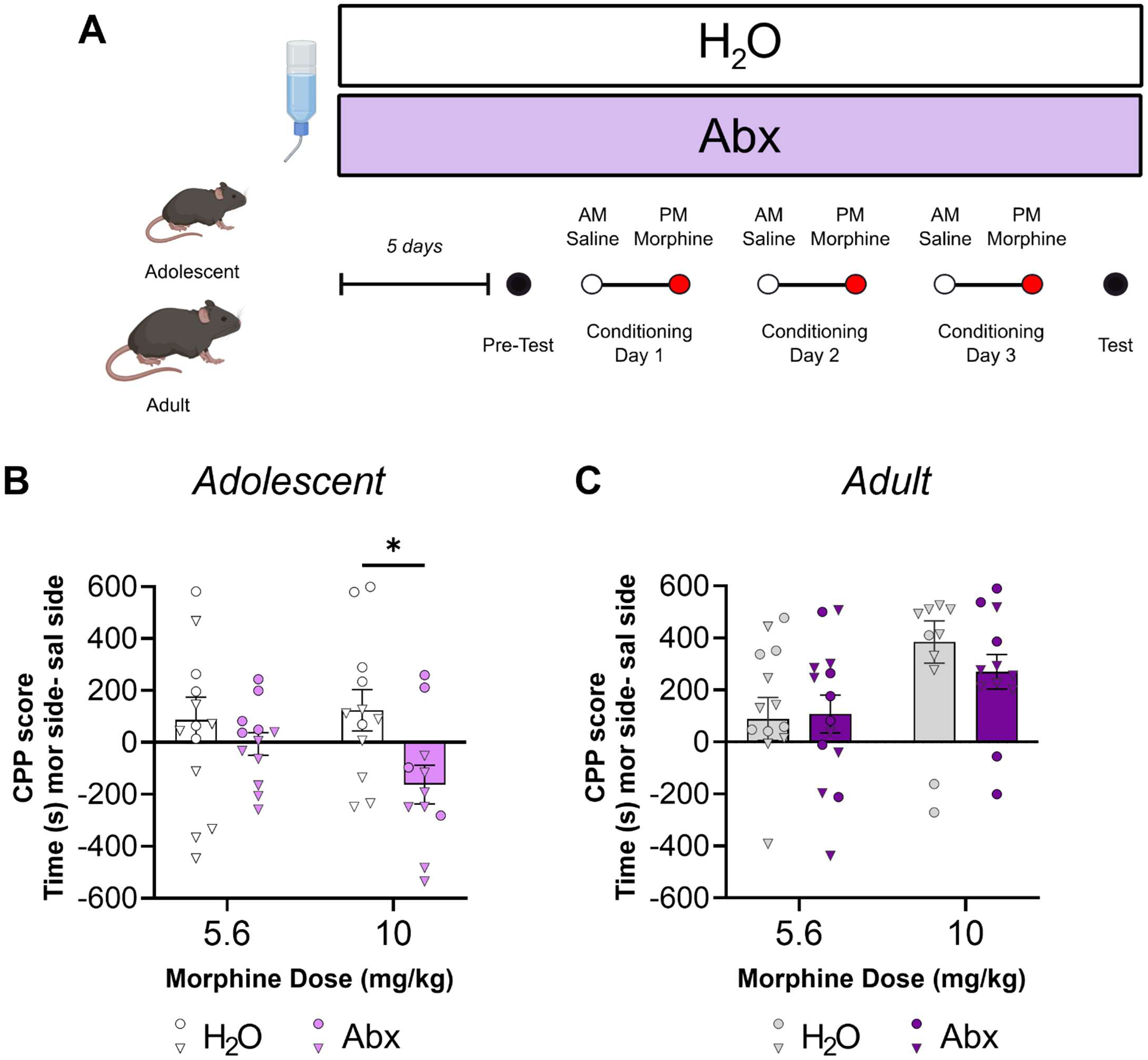
Adolescents display reduced morphine place preference after brief Abx. **(A)** Experimental timeline. **(B)** Morphine conditioned place preference in adolescent mice. There was a main effect of Abx (p = 0.0159), and a significant pairwise comparison between H_2_O and Abx at the 10 mg/kg morphine dose (p = 0.0247), but no other significant effects. **(C)** Morphine conditioned place preference in adults. There was a main effect of morphine dose (p = 0.0044) but no other significant effects in adults, *p < 0.05, males (•), females (▾).

### 3.2. Brief Abx alters microbial diversity in adolescent and adult mice

Given age-specific differences observed in morphine reward, we next determined if Abx exposure altered the microbiome of adolescent and adult mice receiving morphine and if this differed by age. In a separate group of animals that did not undergo behavioral testing, mice received five days of saline or morphine injections (15 mg/kg, s.c.) after 5 days of Abx or H_2_O as described (**Fig. 2A**). Abx consistently decreased the richness of the microbiome in every group tested as measured by Chao1 index (**Fig. 2B & D**; main effect of treatment males: *F*_(2,24)_ = 65.93, *p* < 0.0001 and females: *F*_(2,23)_ = 39.31, *p* < 0.0001; all pairwise comparisons between H_2_O-Sal and Abx-Mor *p* < 0.01). However, morphine alone affected richness in females (**Fig. 2D**; adolescents: *p* = 0.0129 and adults *p* = 0.0014). In contrast to richness, brief Abx only decreased microbiome evenness in adults (**Fig. 2C & E**; Simpson index main effect of treatment males: *F*_(2,24)_ = 11.59, *p* = 0.0003, pairwise comparison *p* < 0.0001 and females: *F*_(2,23)_ = 8.996, *p* = 0.0013, pairwise comparison *p* < 0.0001). Visual examination of beta diversity identified four unique microbiome clusters in male mice. These clusters were separated by age and Abx exposure (**Fig. 2F**). However, females did not show this discrete clustering by age at baseline- the only segregating factor was Abx (**Fig. 2G**).

**Fig. 2.**
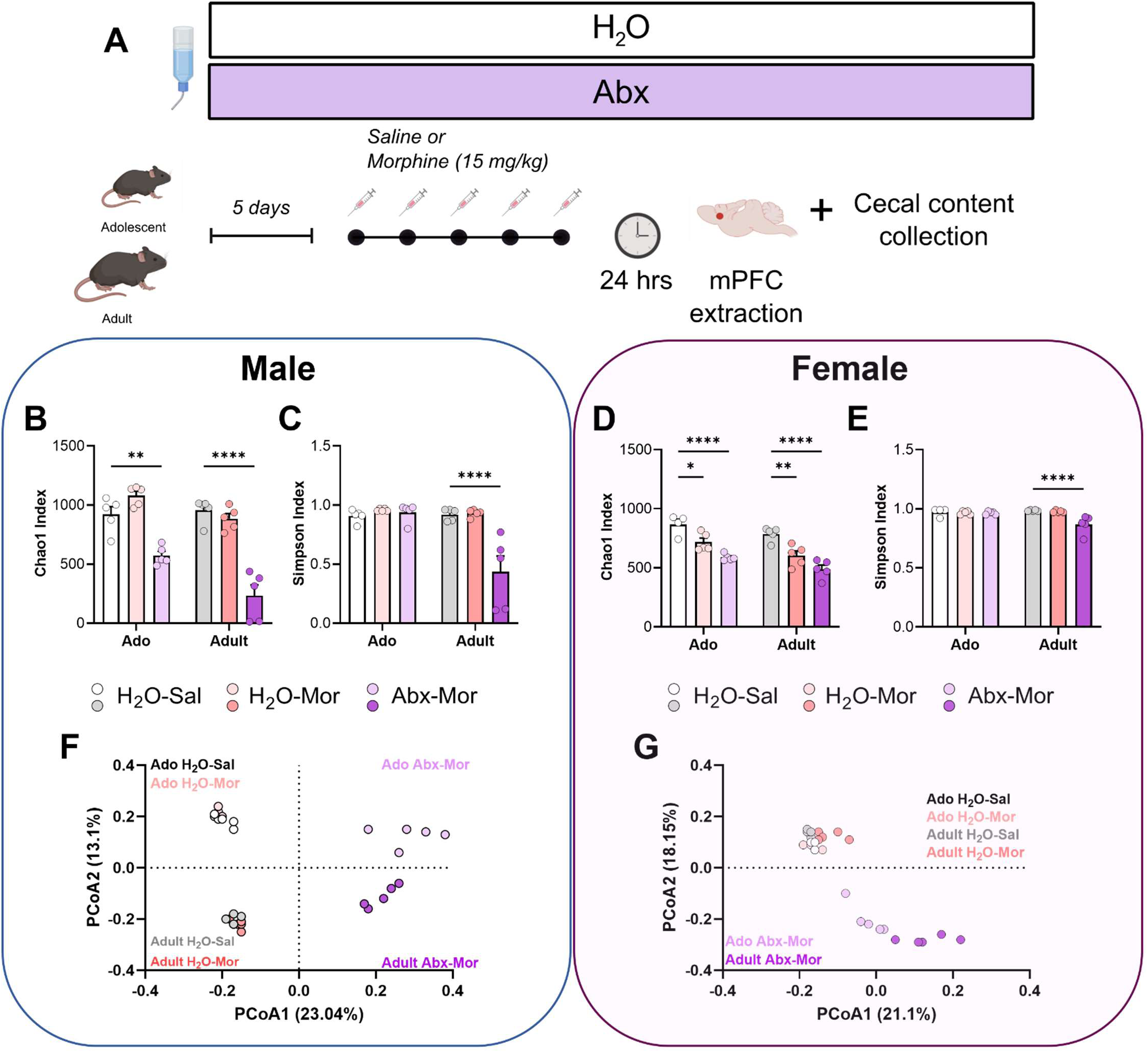
Brief Abx reduces microbiome diversity in morphine-treated mice regardless of age or sex. (A) Experimental timeline. (B & D) Chaol index of species richness in male (B) and female (D) mice. (C & E) Simpson index of species evenness in male **(C)** and female **(E)** mice. **(F & G)** Principal coordinate analysis plots using unweighted Unifrac distance of microbiome samples from male **(F)** and female **(G)** mice. * p < 0.05, ** p < 0.01, **** p < 0.0001.

### 3.3. Gene expression in mPFC differs between adolescent and adult mice

Since brief Abx reduced morphine CPP in adolescent, but not adult mice despite more prominent alterations to microbial diversity in adults, we next wanted to investigate differences in gene expression within the mPFC. Any age-dependent gene expression changes in mPFC could help explain behavioral differences observed in morphine CPP, given the importance of this region in opioid CPP^14^ and its rapid development during adolescence.

We first investigated baseline age differences in gene expression by comparing adolescent H_2_O-Sal to adult H_2_O-Sal in males and females. There were more differentially regulated genes between adolescent and adult males (**Fig. 3B**, 437 downregulated, 555 upregulated) than females (**Fig. 3F**, 23 downregulated, 40 upregulated). Pathway analysis identified several pathways related to synapse pruning in males, such as “neuron projection development” and “synaptic signaling” which were predicted to be downregulated in adolescents (**Fig. 3C**) and “regulation of neuron apoptotic process” which was upregulated (**Fig. 3D**). In contrast, several upregulated terms in adolescent females included those related to myelination, such as “ensheathment of neurons”, “gliogenesis”, “oligodendrocyte differentiation”, and “myelination” (**Fig. 3H**). As expected, age differences in mPFC gene expression seem to reflect known developmental changes that occur during adolescence. To investigate whether morphine or Abx influence the adolescent-specific gene signatures, heatmaps depicting fold-change expression in DEGs from the adolescent gene signature were generated and fold-change of adolescent H_2_O-Mor and adolescent Abx-Mor compared to adult controls of corresponding genes were aligned beneath (**Fig. 3E & I**). In both males and females, the developmental signature seems to weaken over treatments, visualized as fewer genes appearing as dark red or dark blue in H_2_O-Mor and Abx-Mor groups (**Fig. 3E & I**). However, most genes that are up- or downregulated in adolescent H_2_O-Sal are regulated in the same direction or do not differ from adult H_2_O-Sal at all, suggesting that Abx and morphine do not produce opposite effects on gene regulation in these adolescent signature genes.

**Fig. 3.**
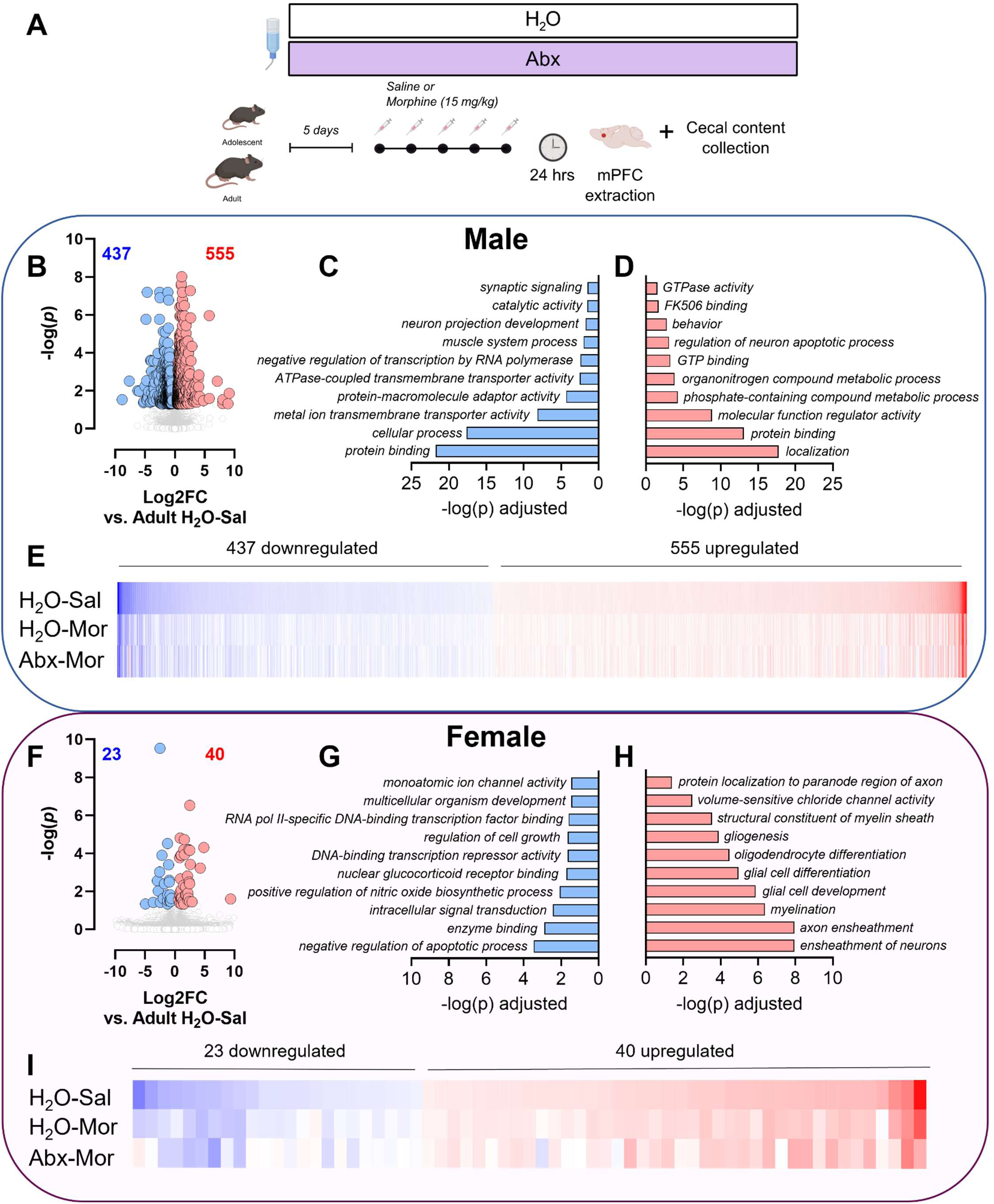
Age differences in gene expression at baseline are only moderately altered by Abx and morphine. **(A)** Experimental timeline. **(B & F)** Volcano plots showing the number of differentially expressed genes (DEG) in male **(B)** and female **(F)** adolescent H_2_O-Sal mice when compared to sex-matched adult H_2_O-Sal mice after FDR correction. Red dots represent significantly upregulated genes, and blue dots represent downregulated genes. **(C & G)** The top ten predicted pathways from male **(C)** and female **(G)** adolescent H_2_O-Sal downregulated DEGs using an FDR corrected -log(p). **(D & H)** The top ten predicted pathways from male **(D)** and female **(H)** adolescent H_2_O-Sal upregulated DEGs using an FDR corrected -log(p). **(E & I)** Heatmap depicting log(2)fold change gene expression of DEGs identified between male **(E)** and female **(I)** adolescent H_2_O-Sal and adult sex-matched H_2_O-Sal in each adolescent treatment group. Top row depicts expression change between adolescent H_2_O-Sal and adult H_2_O-Sal ordered from most downregulated on the left (blue) to the most upregulated on the right (red). The middle and bottom rows reflect log(2)fold change gene expression between adolescent H_2_O-Mor compared to adult H_2_O-Sal (middle) and adolescent Abx-Mor compared to adult H_2_O-Sal (bottom) aligned with the corresponding DEG in the top row between adolescent and adult controls (H_2_O-Sal).

### 3.4. Abx and morphine alter mPFC gene expression differently in adolescents and adults

As we have demonstrated previously in the NAc, Abx during drug exposure drastically enhances the number of differentially regulated genes when compared to drug administration alone^36,38^. In males, Abx-Mor adolescents had 1,170 and adults had 723 DEGs compared to 103 and 1 DEGs in H_2_O-Mor adolescent and adult mice, respectively (**Fig. 4A**). The majority of DEGs in Abx-Mor adolescents were unique to this treatment group, but this was also true for adults (**Fig. 4B**). Pathway analysis of unique adolescent Abx-Mor DEGs identified upregulated pathways relating to transcription and DNA binding, while downregulated pathways varied but included “response to wounding” and “basement membrane”, as well as pathways related to synapses such as “synaptic receptor adaptor activity” and “structural constituent of postsynaptic density” (**Fig. 4C**).

**Fig. 4.**
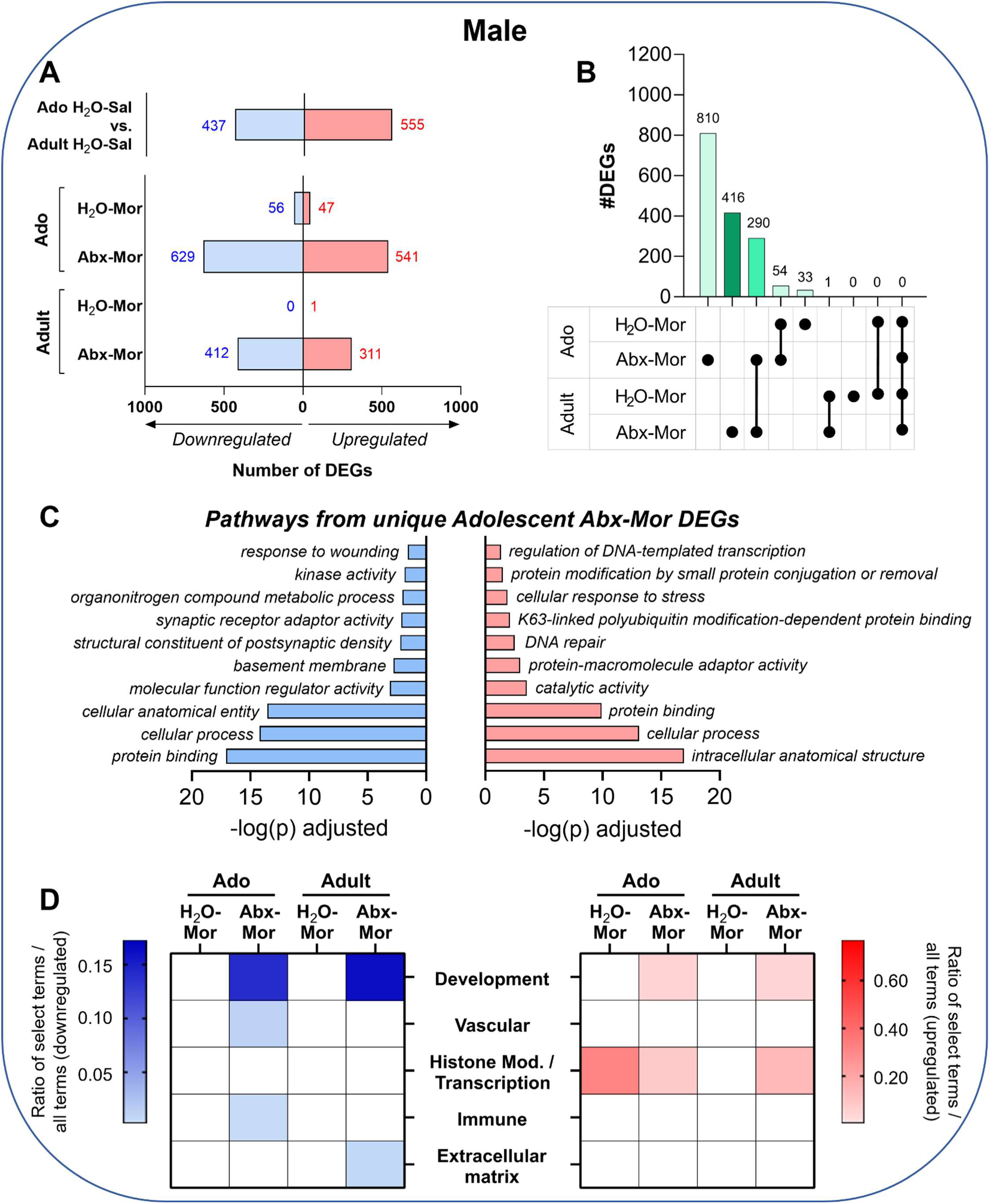
Treatment effects on gene expression differ by age in male mPFC. **(A)** Total number of differentially expressed genes in each treatment group when compared to their age-matched control (H_2_O-Sal). **(B)** Upset plot depicting the total number of unique differentially expressed genes in each treatment group or set of treatment groups. Groups are indicated in the bottom portion of the figure by singular black dots or a series of black dots connected by a vertical line. **(C)** Significant pathways from unique DEGs from adolescent Abx-Mor (compared to adolescent H_2_O-Sal). Pathways from downregulated genes are in blue (left) and pathways from upregulated genes are in red (right). **(D)** Heatmaps depicting the ratio of pathways containing selected keywords to total number of pathways in each comparison. Downregulated terms are in blue (left) and upregulated terms are in red (right).

Next, given the plethora of pathways identified across all pairwise comparisons, we performed an *a priori* keyword search of pathway names. Given the hypothesis that Abx would alter the mPFC transcriptome of adolescents differently than adults, we first searched for pathway terms related to development. Additionally, based on our prior work^36,38^ we searched for terms related to histone modification and transcription. Finally, based on emerging literature, we looked for terms related to vasculature^53^, immune functioning^54,55^, and the extracellular matrix^30,56^. There were several terms relating to development that were both up- and downregulated in male Abx-Mor mice only (**Fig. 4D**). Like what we have observed in the NAc after Abx-Mor^38^, adult Abx-Mor males had a higher ratio of upregulated histone modification and transcription terms than adult H_2_O-Mor. Interestingly, this effect was not present in adolescents, where there were a higher ratio of these terms in H_2_O-Mor mice (**Fig. 4D**). Vascular and immune terms were predicted downregulated in adolescent Abx-Mor and extracellular matrix was predicted to be downregulated in adult Abx-Mor.

As observed in males, female Abx-Mor had the largest number of DEGs- 1,220 DEGs in Abx-Mor adolescents and 305 in Abx-Mor adults compared to 27 and 110 in H_2_O-Mor adolescents and adults (**Fig. 5A**). Similar to males, the adolescent Abx-Mor group had the greatest number of unique DEGs. Pathway analysis of the unique adolescent Abx-Mor DEGs produced predicted downregulated pathways including those related to cell adhesion, such as “regulation of cell-substrate adhesion” and “cell-cell signaling”. However, upregulated pathways were similar to males and included pathways related to transcription such as “DNA binding” and “transcription coactivator activity” (**Fig. 5C**).

**Fig. 5.**
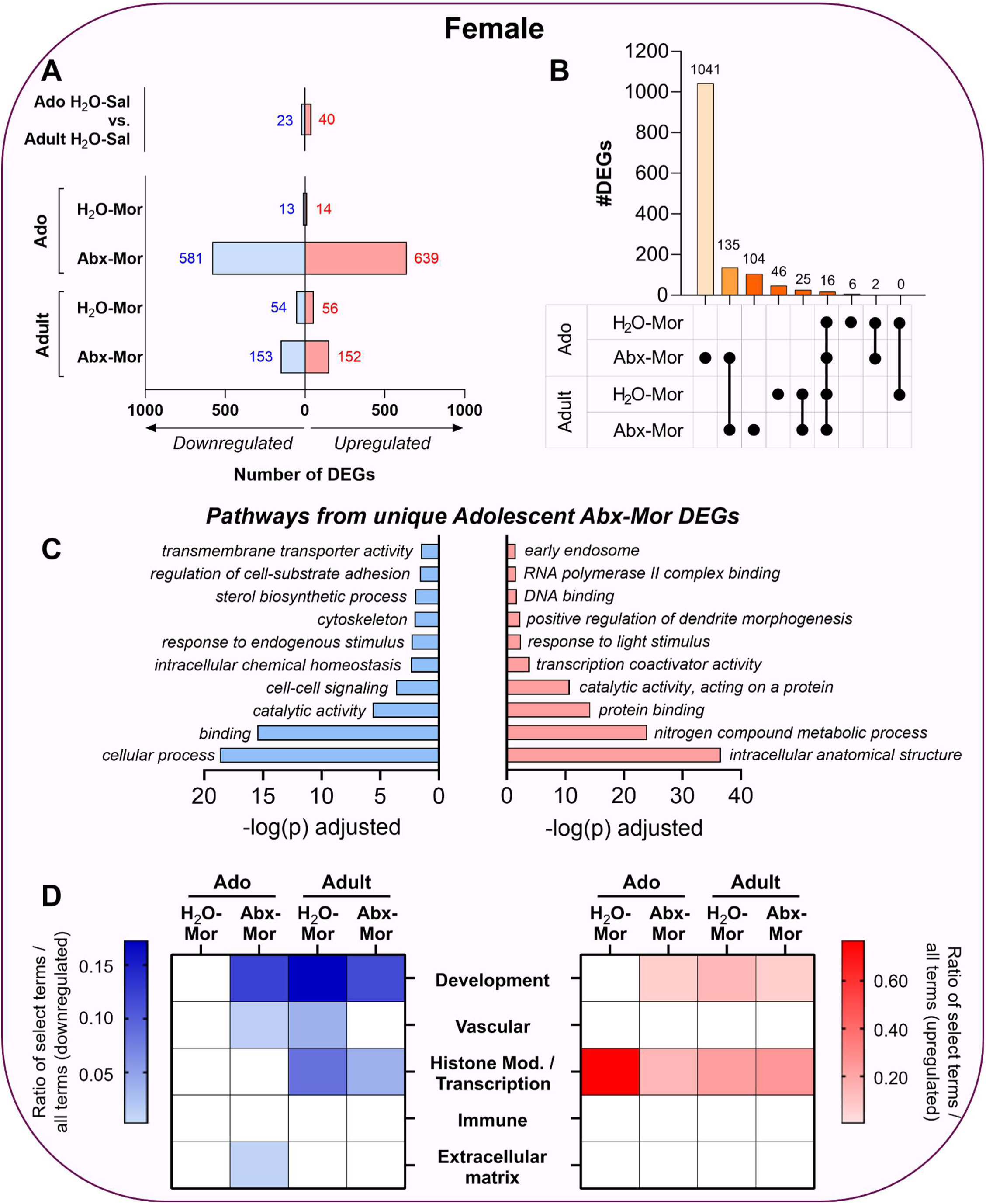
Treatment effects on gene expression differ by age in female mPFC. **(A)** Total number of differentially expressed genes in each treatment group when compared to their age-matched control (H_2_O-Sal). **(B)** Upset plot depicting the total number of unique differentially expressed genes in each treatment group or set of treatment groups. Groups are indicated in the bottom portion of the figure by singular black dots or a series of black dots connected by a vertical line. **(C)** Significant pathways from unique DEGs from adolescent Abx-Mor (compared to adolescent H_2_O-Sal). Pathways from downregulated genes are in blue (left) and pathways from upregulated genes are in red (right). **(D)** Heatmaps depicting the ratio of pathways containing selected keywords to total number of pathways in each comparison. Downregulated terms are in blue (left) and upregulated terms are in red (right).

In females, development terms were up- and downregulated in all groups except adolescent H_2_O-Mor (**Fig. 5D**). Additionally, all morphine treated groups had several histone modification terms that were upregulated, as well as some that were downregulated in adults (**Fig. 5D**). Vascular terms were found to be predicted downregulated in adolescent Abx-Mor and adult H_2_O-Mor; extracellular matrix was predicted to be downregulated adolescent Abx-Mor only.

Finally, we examined global patterns of gene expression change using a threshold free analysis. Weighted gene coexpression network analysis (WGCNA) uses a data-driven approach to identify groups of genes that change their expression in the same direction across all samples or within select groups. Notably, WGCNA was not performed on male subjects as no detected network met our inclusion criteria previously outlined (i.e. networks with > 40 genes). In females, of 80 total modules, 36 produced significant correlations in at least one treatment group (**Fig. 6A**). Of these, we prioritized modules with differential directional regulation between adolescent H_2_O-Mor and adolescent Abx-Mor (black module) and the two modules with the most associated pathways that were significantly regulated in adolescent (blue) or adult (light green) Abx-Mor only.

**Fig. 6.**
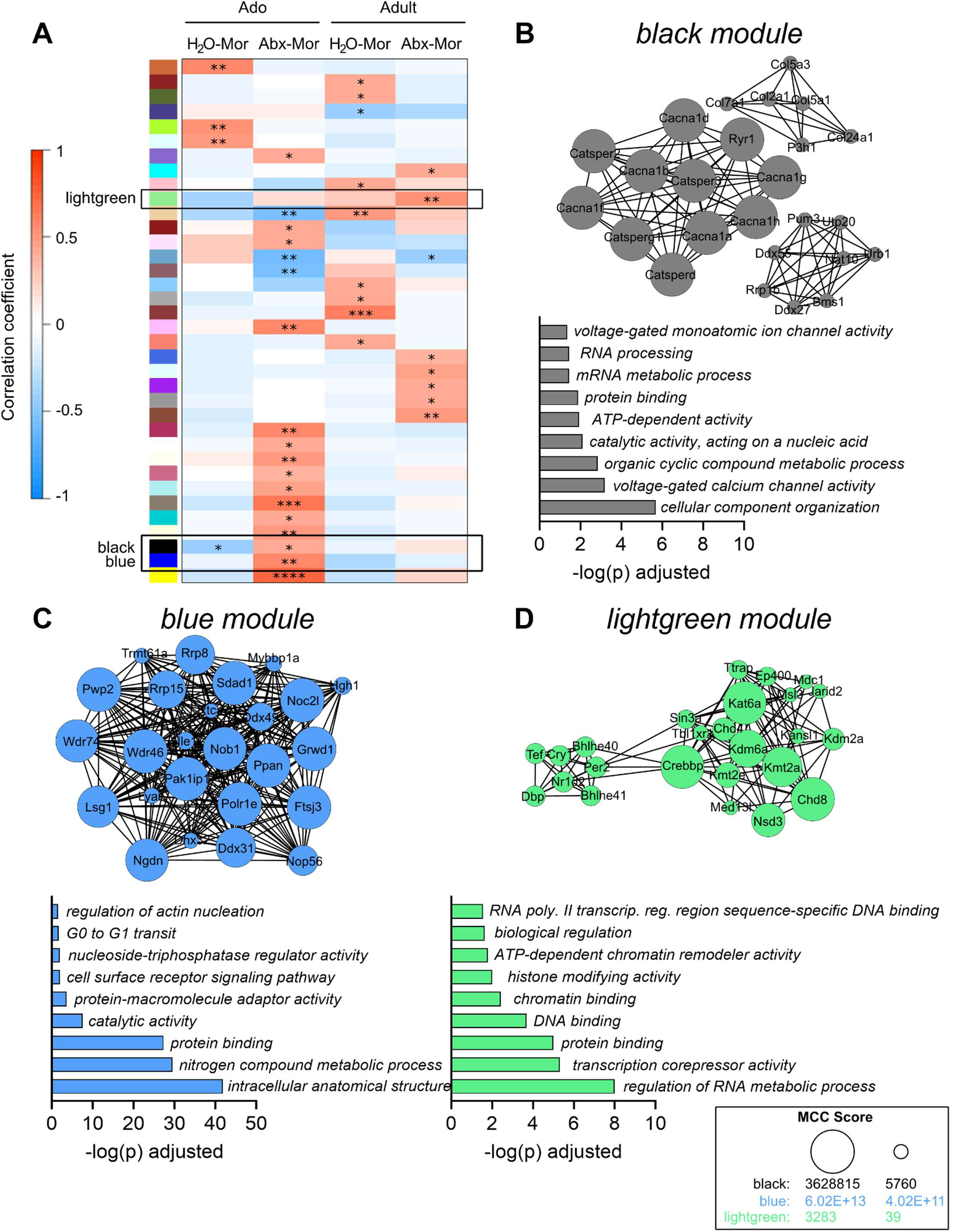
WGCNA identified unique expression modules in female adolescents and adults. **(A)** Correlation heatmap of selected modules. The module color is indicated to the left of the correlation plot. Positive correlations are shown in red and negative correlations are in blue. Asterisks indicate significant correlations (* p < 0.05, ** *p* < 0.01, *** *p* < 0.001, **** *p* < 0.0001). Only modules that were significant in at least one treatment group are shown. Adolescent and adult control groups also not shown for clarity. The modules surrounded by black boxes were selected for further analysis (lightgreen, black, and blue). **(B)** The top 25 nodes in the black module (top). The top 9 pathways from all nodes in the black module (bottom). **(C)** The top 25 nodes in the blue module (top). The top 9 pathways from all nodes in the blue module (bottom). **(D)** The top 25 nodes in the green module (top). The top 9 pathways from all nodes in the green module (bottom). Node size adjusted separately for each module using MCC score (legend on bottom right).

The black module, that was significantly negatively correlated in adolescent H_2_O-Mor and positively correlated in adolescent Abx-Mor, contained 3 clusters of genes-notably one encoding calcium channel subunits (*Cacna1a, Cacna1b, Cacna1d, Cacna1f, Cacna1g, Cacna1h*) and another encoding RNA helicases (e.g. *Ddx27, Ddx55*). Overall, pathways indicated that nodes in this module are involved in calcium signaling (“voltage-gated calcium channel activity” and “voltage-gated monoatomic ion channel activity”) and RNA metabolism (“RNA processing”, “mRNA metabolic process”, and “catalytic activity, acting on a nucleic acid”). The blue module had the most genes of any modules studied with the highest MCC scores overall. This module was significantly positively correlated in adolescent Abx-Mor only. Pathways were varied, but contained “cell surface receptor signaling pathway”, “regulation of actin nucleation”, and “G0 to G1 transit”. Finally, the light green module (significant in adult Abx-Mor only), contained many histone modifiers such as lysine acetyltransferases (e.g. *Kat6a*), lysine demethylases (e.g. *Kdm6a* and *Kdm2a*), and lysine methyltransferases (e.g. *Kmt2a, Kmt2e*). This is reflected in the associated pathways “ATP-dependent chromatin remodeler activity”, “histone modifying activity”, “chromatin binding”, and “DNA binding”.

## 4. Discussion

This collection of studies identified an age-specific effect of brief Abx on morphine reward and gene expression after morphine in the mPFC of mice. In contrast to our past study that found reduced place preference in adult male mice^38^, here we found that adolescents, but not adults, had reduced morphine place preference after Abx. In the current study, we employed a shorter exposure period of Abx (5 days instead of 14 days), and this experimental difference likely explains the discrepancy with our earlier work. Our work here suggests that adolescents are more sensitive to the behavioral effects of microbiome disruption since adults showed reduced CPP after 14 days Abx pretreatment but adolescents demonstrated that same effect after only 5 days. This work adds support to previous studies demonstrating that adolescent microbiome disturbance, even if brief, can have marked effects on behavior^23,30,31^.

Since adolescents are undergoing many physiological changes, identifying the mechanism underlying the age effect of microbiome disturbance on morphine reward is challenging. Given that both the mPFC and the microbiome are undergoing significant changes during adolescence^15–20^, we investigated how brief Abx, together with morphine, changes the microbiome and gene expression within the mPFC. Sequencing of the microbiome confirmed underlying age differences in the microbiome of male mice (but not females) and found that brief Abx changed the microbiome in all groups of mice tested. However, the overall effects of Abx were mild compared to that observed in prior studies^36–38^. As previously seen, Abx decreased species richness or abundance (measured as Chao1 index) in all groups. Interestingly, species abundance was also decreased in H_2_O-Mor females at both ages, an effect we have not observed in adult male animals after opioids^37,38^. However, species evenness was only reduced after Abx in adult mice, potentially suggesting that adults are more sensitive to the microbiome-altering effects of Abx. This is contrary to behavior, where adolescents are more sensitive to the reward-reducing effects of Abx. Taken together, the effect of brief Abx on morphine CPP is unlikely to be explained solely by age differences in the extent of microbiome disruption produced by Abx.

To further examine the biological locus of our behavioral results, this study measured gene expression changes in the mPFC of adolescents and adults after morphine with and without microbiome disruption. First, we identified an adolescent-specific gene expression signature by comparing gene expression between adolescent and adult H_2_O-Sal controls. Unsurprisingly, genes involved in synapse pruning and myelination were differentially regulated between adolescents and adults. However, there was a sex effect-males showed downregulation of synapse related terms and females showed upregulation of myelination-related terms. While both increases in myelination and decreases in synapse density occur during adolescence, these sex dependent DEG changes suggest they might occur differently in males and females or might occur at slightly different ages in males and females.

As in our previous dataset^38^, Abx-Mor mice expressed more DEGs than age- and sex-matched H_2_O-Mor groups. Overall, adolescent Abx-Mor expressed the most total DEGs and the most unique DEGs compared to all other groups. Despite differences in total numbers of DEGs, pathway analysis indicated that all Abx-Mor mice had upregulation of pathways affecting gene transcription and histone modification, an effect we have seen previously in the NAc^38^. Given that so many of the Abx-Mor DEGs between adolescents and adults do not overlap, it is possible that the transcription factors and chromatin modifiers within these pathways are not the same in every Abx-Mor group. Unlike in adult males that only had altered histone modification pathways in Abx-Mor groups, females and adolescent male H_2_O-Mor mice also had upregulated histone modification pathways as well.

We also identified several development pathways that were predicted to be up and downregulated in many groups-not just adolescents. In fact, every Abx-Mor group had up- and downregulated terms related to development. This is not true of H_2_O-Mor groups where only adults had differences in development terms and adolescents had none. Surprisingly, we found no overall pattern of vascular, extracellular matrix, or immune-related terms across comparisons. These terms appeared infrequently, although when they were present, they were always downregulated and almost exclusively in Abx-Mor groups. Despite their sparse appearance in our data, the mechanisms described in these rare terms could be important in understanding how adolescents and adults differ in their response to Abx and morphine.

Finally, threshold free analysis using WGCNA identified several noteworthy modules in female mice. Analysis of the black module, which was positively correlated in adolescent Abx-Mor females and negatively correlated in adolescent H_2_O-Mor, identified calcium channels and genes involved in RNA metabolism as separate clusters of nodes. Currently, any relationship between these disparate groups of genes has not been explored, especially in the context of cortical functioning or responses to opioids. We also analyzed the light green module, which was upregulated in adult female Abx-Mor only. This module contained several different classes of histone modifiers, in line with pathway analysis in the female Abx-Mor mice and our prior work in adult males^36,38^.

This study examined the behavioral and transcriptional response to brief Abx and morphine in adolescent and adult mice, where we found an adolescent-specific reduction in morphine CPP that corresponded to unique changes in gene expression within mPFC. While much work is needed to determine the causal relationship between transcriptional changes in cortex to behavior, this study provides many promising avenues to investigate including age-specific histone modifications and alterations to the extracellular matrix and their role in opioid reward. Understanding how adolescents respond to drugs of abuse and understanding how environmental stimuli influence these responses is necessary to develop policies to reduce adolescent drug use.

## Acknowledgements

Funding: This work was supported by the National Institutes of Health [R01DA056592 to DDK and K01DA050906 to RSH].

